# DNase treatment improves viral enrichment in agricultural soil viromes

**DOI:** 10.1101/2021.06.01.446688

**Authors:** Jackson W. Sorensen, Laura A. Zinke, Anneliek M. ter Horst, Christian Santos-Medellin, Alena Schroeder, Joanne B. Emerson

## Abstract

The small genomes of most viruses make it difficult to fully capture viral diversity in metagenomes dominated by DNA from cellular organisms. Viral size-fraction metagenomics (viromics) protocols facilitate enrichment of viral DNA from environmental samples, and these protocols typically include a DNase treatment of the post-0.2 μm viromic fraction to remove contaminating free DNA prior to virion lysis. However, DNase may also remove desirable viral genomic DNA (*e.g*., contained in virions compromised due to frozen storage or laboratory processing), suggesting that DNase-untreated viromes might be useful in some cases. In order to understand how virome preparation with and without DNase treatment influences the resultant data, here we compared 15 soil viromes (7 DNase-treated, 8 untreated) from 8 samples collected from agricultural fields prior to tomato planting. DNase-treated viromes yielded significantly more assembled viral contigs, contained significantly less non-viral microbial DNA, and recovered more viral populations (vOTUs) through read mapping. However, DNase-treated and untreated viromes were statistically indistinguishable, in terms of ecological patterns across viral communities. Although results suggest that DNase treatment is preferable where possible, in comparison to previously reported total metagenomes from the same samples, both DNase-treated and untreated viromes were significantly enriched in viral signatures by all metrics compared, including a ~225 times greater proportion of viral reads in untreated viromes compared to total metagenomes. Thus, even without DNase treatment, viromics was preferable to total metagenomics for capturing viral diversity in these soils, suggesting that preparation of DNase-untreated viromes can be worthwhile when DNase treatment is not possible.

**Importance:** Viromics is becoming an increasingly popular method for characterizing soil viral communities. DNase treatment of the viral size fraction prior to DNA extraction is meant to reduce contaminating free DNA and is a common step within viromics protocols to ensure sequences are of viral origin. However, some samples may not be amendable to DNase treatment due to viral particles being compromised either in storage (i.e. frozen) or during other sample processing. To date, the effect of DNase treatment on the recovery of viruses and downstream ecological interpretations of soil viral communities is not thoroughly understood. This work sheds light on these questions and indicates that while DNase treatment of soil viromes improves recovery of viral populations, this improvement is modest in comparison to the gains made by viromics over total soil metagenomics. Further, DNase treatment may not be necessary to observe the ecological patterns structuring soil viral communities.

## Introduction

Viruses infect all three domains of life and play key roles not only in human health but also in agriculture and global nutrient cycling (1–5). They are important in oceanic food webs, and our understanding of their role in soils is growing rapidly (2, 6–14). Viral abundances are estimated to range from 10^7^ to 10^10^ virions per gram in soil (6, 11), and measurements from transmission electron microscopy suggest that up to 28% of microbial cells in soil are actively infected by viruses (15–17). Through metagenomic approaches, soil viral populations have been implicated in soil carbon cycling and microbial community dynamics in changing environments, including in thawing permafrost and other peatlands (18–20).

The study of soil viral communities has lagged behind analogous efforts in marine systems, in part because the complex and heterogeneous nature of soil presents unique challenges for recovering viral DNA (2, 10, 11, 14, 21). Although marine viral ecology has benefitted from a viromics approach, in which purified, concentrated viral particles are subjected to DNA extraction and metagenomic sequencing (12, 22, 23), most recent soil viral ecological studies have focused on recovering viral signatures from total soil metagenomes (10, 19, 20, 24). Bioinformatic advances in viral contig identification (*e.g*., through recognition of viral hallmark genes and other viral sequence signatures) (25–28) and efforts to compile viral reference databases that include partial and putative viral genomes (1, 19, 29) have improved our ability to recognize viral genomic sequences in soil metagenomes. However, despite these advances, our ability to catalog soil viral diversity is still largely gated by the low prevalence of viral DNA in total soil metagenomes, which tend to be dominated by bacterial and archaeal sequences (10, 19, 30).

Fortunately, viral size-fractionation protocols (*e.g*., passage of a sample through a 0.2 μm filter to remove most cells), initially used in marine and other aquatic systems (12, 22, 23, 31–34), have also been applied to soil (11, 35), and recent data suggest that these protocols can enrich the viral signal in sequencing data (10, 19, 21, 30). Through iterative steps of mechanical and/or chemical desorption and centrifugation, virus-sized particles are separated from the soil matrix and from microbial cells, and DNA can then be directly extracted and sequenced from this viral size fraction to generate a shotgun metagenome, known as a virome (11, 18, 19, 30, 36, 37). Our group has shown that this approach can greatly increase both the number of viral populations and the proportion of viral DNA in the produced sequencing data from soil viromes, compared to total metagenomes (19, 30). For example, in agricultural soils, on average 30 times more contigs were identified as viral and 585 times more reads were recruited to viral genomes in viromes, compared to total metagenomes from the same samples (30).

A common step in laboratory viromics protocols is treatment with DNase after the post-0.2 μm (viral) fraction has been purified and enriched but before DNA extraction (11, 21, 30, 37). Under the assumption that most viral particles (virions) remain intact with their genomic contents protected at this stage, a DNase treatment is meant to reduce the amount of extracellular and/or free, “relic” DNA (38) that may have been co-enriched with the virions. The amount of “relic” DNA in a given soil sample presumably varies widely, depending on the soil, and the amount recovered in a given metagenomic or viromic library will also depend on the laboratory procedure(s) used to prepare the DNA (38–40). Estimates of relic DNA in soil vary (38, 39), but one study suggested that, on average, 40.7% of soil 16S rRNA gene amplicon sequences are relic (38). One meta-analysis of viromes (predominantly from freshwater, saline, and human gut environments, with none from soils) determined that a range of 0.2 % to 40.3 % of viromic reads were mapped to non-viral microbial genomes, suggesting the potential for substantial non-viral DNA contamination in some cases (41). However, the amount of free DNA contamination in soil viromes and the potential impact of this DNA on downstream analyses has yet to be thoroughly considered.

Although these prior results would suggest that DNase treatment is an important step in the process of preparing a virome, virions themselves can be compromised prior to DNA extraction, such that DNase treatment of these compromised virions may remove the very viral genomic DNA that was meant to be enriched. Virions can be compromised both naturally through degradation in the environment and potentially during sample collection, transportation, storage, and/or laboratory processing (42–44). In some cases, particularly if the virions were compromised after removal from the field, it may be desirable to recover DNA from these compromised virions. The successful enrichment of viral DNA via viromics without DNase treatment has been previously observed, for example, from a hypersaline lake and from soil (peat) samples stored frozen (19, 34). This suggests that, in cases in which DNase treatment of a virome is not possible due to loss of viral DNA, preparation of a virome that has not undergone DNase treatment may still be worthwhile. However, direct comparisons of DNase-treated and untreated viromes from the same samples have not been made in soil (or any other environment, to our knowledge), nor have these two types of viromes been placed in the context of recoverable viral sequences from total metagenomes.

Here we sought to better understand the differences between soil viromes prepared with and without DNase treatment, in order to more thoroughly evaluate the utility of non-DNase-treated soil viromes (hereafter, untreated viromes). Considering 15 viromes (7 DNase-treated (previously reported, with one having failed at the library construction step (30)), and 8 untreated (new in this study)) from 8 agricultural soil samples, this study compares overall sequence complexity, assembly success, proportions of recoverable viral contigs, the percentage of viral reads, viral taxonomic diversity, and the downstream ecological interpretations that would be derived from these two treatments. We hypothesized that treatment with DNase would increase the recovery of viral contigs by decreasing the overall sequence complexity and improving assembly and, therefore, that DNase treatment would be preferable, where possible. We also suspected that the overall patterns of viral community beta-diversity across samples would not be significantly influenced by DNase treatment and that untreated viromes would yield substantially more recognizable viral sequences than the total metagenomes that were previously sequenced from these same samples (30).

## Materials and Methods

### Sample collection and soil processing

Our sampling design and soil collection have been described previously (30). Briefly, eight agricultural tomato plots near the UC Davis campus (38°32’08”N, 121°46’22” W) were sampled on April 23^rd^ of 2018. Each of the plots had been treated with one of four biochar amendments (650 °C pyrolyzed pine feedstock, 650 °C pyrolyzed coconut shell, 800 °C pyrolyzed almond shell, or no biochar control) on November 8^th^, 2017 as part of an ongoing study to investigate the impact of biochar treatment on agricultural production (Table S1). Tomato seedlings had not yet been planted at the time of sampling (the field was fallow). The top 30 cm of soil was collected using a 2.5 cm diameter probe, and a total of 8 probe cores per plot were combined into a single sterile bag per sample and transported on ice to the laboratory, where each sample was sieved through an 8 mm mesh.

### Viral purification and DNA extraction for viromics

The eight DNase-treated viromes were prepared as previously described (30), and in the current study, the same soils were also prepared without DNase treatment, for a total of 16 samples. Laboratory processing for all samples was the same up to the DNase treatment step. Briefly, viromes were generated for each sample from 50 g of fresh soil separated into two 50 mL conical tubes, following a previous protocol (11) with slight modifications (30). To each of the two tubes per sample, 37.5 mL of 0.02 μm filtered AKC’ extraction butter (10% PBS, 10 g/L potassium citrate, 1.44 g/L Na_2_HPO_4_, 0.24 g/L KH_2_PO_4_, 36.97 g/L MgSO_4_) (37) was added. Tubes were briefly vortexed to homogenize the soil slurry and then shaken at 400 RPM for 15 minutes on an orbital shaker. Subsequently, each tube was vortexed for an additional 3 minutes before undergoing centrifugation at 4,700 x g for 15 minutes to pellet the soil. The two supernatants from the same sample were then filtered through a 0.22 μm polyethersulfone filter to remove most cells and combined into a 70 mL polycarbonate ultracentrifuge tube, which was centrifuged for three hours at 4 °C and 32,000 x g to pellet viral particles. Taking care not to disturb the pellet, the supernatant was discarded and the viral pellet resuspended in 200 μL of ultrapure water. The eight untreated samples (no DNase treatment) proceeded directly to DNA extraction at this point. To the eight samples designated for DNase treatment, as previously described (30), 30 units of RQ1 RNase-free DNase and 30 μL of 10X DNase buffer (Promega Corp., Madison, WI, USA) were added, and samples were incubated at room temperature for two hours before stopping the reaction with 30 μL of the DNase stop solution (Promega Corp. Madison, WI, USA). The eight DNase-treated samples underwent DNA extraction at this point. For both DNase-treated and untreated viromes, DNA was extracted using the DNeasy PowerSoil Kit (Qiagen, Hilden, Germany), according to the manufacturer’s instructions with slight modifications, as previously described (30).

### Library construction and sequencing

Libraries were constructed and sequenced by the DNA Technologies and Expression Analysis Core at the UC Davis Genome Center. The DNA Hyper Prep library kit (Kapa Biosystems-Roche, Basel, Switzerland) was used for all libraries. A single lane of Illumina HiSeq 4000 paired-end 150 bp sequencing was used to generate all of the sequencing data with a targeted sequencing depth of 4 Gbp per sample. Raw sequences can be found under the BioProject accession PRJNA646773.

### Sequence processing, assembly, and identifying viral contigs

All viromes were bioiniformatically processed from raw sequencing data (*i.e.*, those that were previously reported were re-processed here). Sequencing reads were quality-trimmed and primers removed using Trimmomatic (45). MEGAHIT was used with the “meta” preset to individually assemble each virome, using the paired quality-trimmed reads and a minimum contig length of 10 kbp (46). Analyses of overall assembly statistics (Figure 1) were performed on these data. Putative viral contigs were then identified using VirSorter in decontamination mode, retaining any contigs that were assigned as higher confidence categories (1, 2, 4, or 5), in accordance with established recommendations (10, 18, 25, 28). Analyses considering viral contigs not yet dereplicated into populations (*i.e.*, Figures 2A-C) were performed on these data.

**Figure 1.**
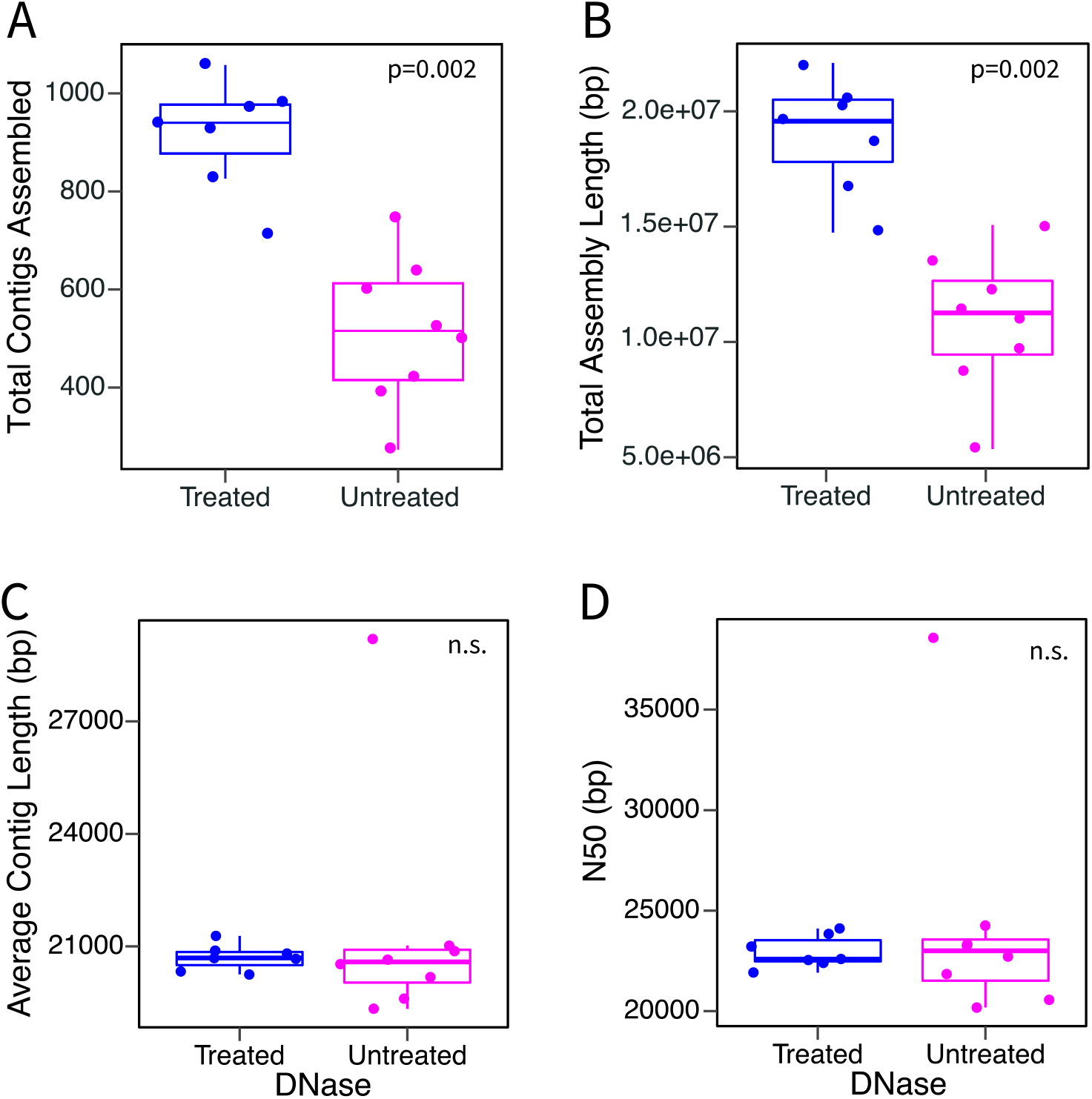
Assembly comparisons of DNase-treated and untreated viromes. Each point is one virome, with comparisons according to: (A) total contigs assembled, (B) total assembly length, (C) average contig length, and (D) N50 (contig length where half the assembly length is represented in longer contigs and half in shorter contigs). Boxes show the interquartile range and median value. Whiskers extend to the furthest non-outlying datapoint. P-values show the significance of Kruskal Wallis tests between DNase-treated (n=7) and untreated (n=8) samples. Insignificant results (p-values >0.05) are listed as n.s.

**Figure 2.**
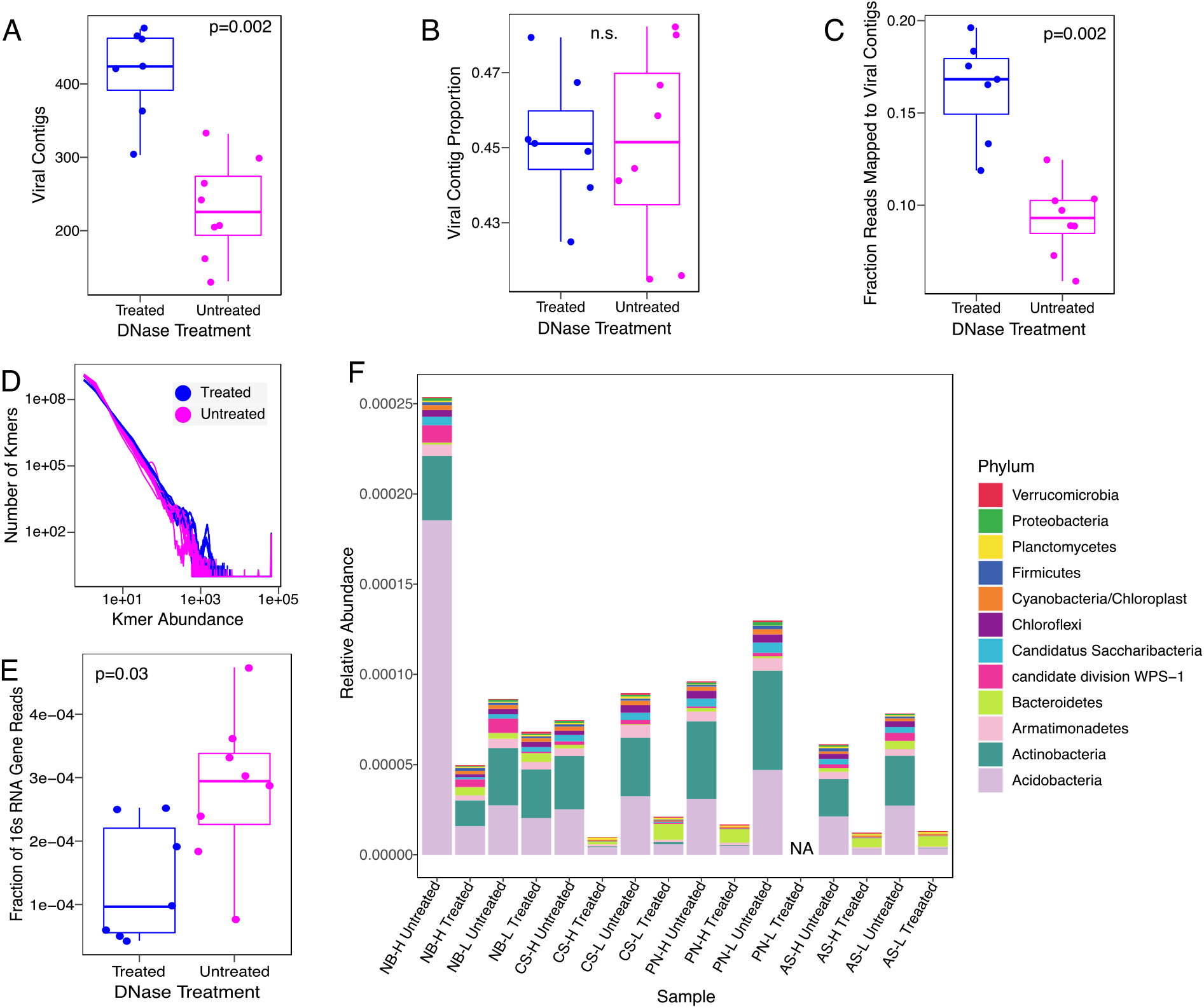
Differences in sequencing content between DNase-treated and untreated viromes. (A) Number of VirSorter identified viral contigs assembled per virome and (B) their proportion of the total number of contigs per virome. (C) The proportion of reads from each sample that mapped to VirSorter identified viral contigs. (D) Frequency plot of kmers showing kmer abundance on the x axis, and the number of kmers with that abundance on the y-axis; each line is one virome. (E) Proportion of reads that contain partial 16S rRNA gene sequences as identified via SortMeRNA. (F) Relative abundances of the top 12 most abundant phyla according to partial 16S rRNA gene sequences. The y-axis displays the number of reads containing 16S rRNA gene fragments from each of the top 12 phyla as a proportion of the total number of quality-trimmed reads in each virome. DNase treated and untreated viromes from the same plot are placed next to each other for ease of comparison. NA indicates no data. For all boxplots (panels A, B, C, and E), boxes show the interquartile range and median value with whiskers extending to the furthest non-outlying datapoint, and P-values show the significance of Kruskal Wallis tests between DNase-treated (n=7) and untreated (n=8) viromes. Insignificant results (p-values >0.05) are displayed as n.s. (not significant).

### Kmer analyses of virome sequence complexity

Kmer counting was performed using the khmer software package version 2.1.1 (47). Reads were first k-mer error trimmed using the command “trim-low-abund.py” before k-mers of size 31 were counted in each virome using the script “load-into-counting.py”.

### Taxonomic identification of bacterial and archaeal 16S rRNA gene content in viromes

SortMeRNA was used with its internal SILVA bacterial and archaeal 16S rRNA gene databases (version V119) to identify partial 16S rRNA gene sequences present in the reads from each virome (48, 49). Reads found to contain partial 16S rRNA gene sequences were then classified using the Ribosomal Database Project classifier trained with the RDP training set and a confidence cutoff of 0.8 (50). Classifications were collapsed at the phylum level to create a phylum-by-sample table in order to investigate changes in the relative abundances of phyla across DNase treatments.

### Viral population (vOTU) identification, read mapping, and vOTU detection criteria for ecological analyses

VirSorter-identified viral contigs (described above) were dereplicated through clustering, using the ‘psi-cd-hit.pl’ command of CDHIT (51) with a minimum alignment length equal to 85% of the smaller contig and minimum percent identity equal to 95%, in accordance with best practices for identifying viral populations (vOTUs) (37). The resulting representative seed contig sequences from each cluster were then used as our set (“database”) of vOTUs for further analysis. vOTU representative seed sequences were annotated using prodigal (52) and then grouped into viral clusters (VCs) and taxonomically identified using vConTACT2 with its “ProkaryoticViralRefSeq85-Merged” database (53).

In order to perform community ecological analyses, the relative abundances of each vOTU in each sample were assessed by read mapping to the reference database of vOTUs. Specifically, quality-trimmed reads were mapped to the database of vOTUs at a minimum identity of 90% using BBMap (54). The resulting SAM files were then converted into sorted and indexed BAM files using samtools (55). The trimmed pileup coverage and read count abundance of each vOTU were calculated using BamM parse to generate tables of vOTU abundances (average coverage depth) in each sample (56). We used bedtools to calculate the per-base coverage for each vOTU in each sample, requiring that >75% of the vOTU contig length be covered by at least one read for detection in a given virome (also known as “breadth”) (57, 58). The vOTU coverage tables generated to this point were considered in analyses with “relaxed” detection criteria, meaning that reads mapped to vOTUs assembled from any sample were included. For analyses using “stringent” detection criteria, we also required that, for a given vOTU to be considered detected in a virome, an assembled contig from that same virome and/or another virome within the same DNase treatment group had to be in the same >=95% nucleotide identity vOTU cluster. In other words, that same vOTU (viral “species”) must have been assembled from a virome in the same treatment group, mimicking a condition in which only that treatment had been performed and thus only vOTUs from that treatment would be in the reference database for read mapping. The resulting vOTU coverage tables were used for downstream ecological and statistical analyses.

### Ecological and statistical analyses

After generating the vOTU tables, all ecological and statistical analyses were performed in R (59). The vegan package was used to calculate Bray-Curtis dissimilarities (function vegdist) using vOTU relative abundances, perform PERMANOVA (function adonis), and correlate matrices (Mantel tests, function mantel) (60). Kruskal.test from the stats package was used to perform the Kruskal-Wallis rank sum test. Boxplots were constructed using ggplot2 and tanglegrams using the dendextend package (61, 62). For both Mantel tests and tanglegram analyses comparing DNase treated and untreated viromes, the untreated virome from plot PN-L was dropped from the analysis because its paired DNase-treated virome failed at the library construction step.

### Data Availability

All viromes analyzed and presented in the current study have been deposited in the NCBI SRA under BioProject PRJNA646773.

## Results

### Comparison of metagenomic assembly success from DNase-treated and untreated viromes

We sampled eight agricultural plots that had been treated with four different biochar amendments (30) and generated two viromes (one treated with DNase and one untreated) from each sample. The DNase-treated viromes were part of a prior study (30) and the untreated viromes are new here. These 16 viromes were sequenced to a depth of 4 Gbp (range 3.65-4.53 Gbp), apart from a single DNase-treated virome from which library construction failed, as previously described (Table S2) (30). Despite equimolar DNA contributions to the sequenced pool of libraries, untreated samples recovered a greater number of sequencing reads compared to their DNase-treated counterparts (Kruskal-Wallis p= 0.02, untreated median = 28,008,452, DNase-treated median = 26,847,586, Supplementary Figure 1). However, after quality filtering, there was no significant difference in the number of reads between treatment types (Kruskal-Wallis p = 0.08). Overall, DNase-treated viromes assembled into significantly more contigs (average 917 DNase-treated contigs, 513 untreated, Kruskal-Wallis p = 0.002) and had a longer total assembly length than their paired untreated viromes (Figure 1). However, the average contig lengths and N50s (*i.e.* the shortest contig length where half the assembly length is represented in longer contigs and half in shorter contigs) were statistically indistinguishable between the two treatments (Figure 1, Table S3).

### Sequence complexity and proportion of cellular organism-derived reads in DNase-treated compared to untreated viromes

We suspected that decreased sequence complexity in DNase-treated viromes contributed to the observed significant improvement in assembly, presumably due to the degradation of “free” DNA (*e.g*., from bacteria and archaea, as opposed to viruses). We tested this in two ways: first by comparing the k-mer complexity between the two approaches, and second by comparing the 16S rRNA gene recovery. DNase-treated viromes tended to have more abundant k-mers and fewer singleton k-mers than their untreated counterparts (Figure 2D), and DNase-treated viromes had significantly fewer total k-mers per sample (Kruskal-Wallis p=0.002). We next asked whether the reduced complexity of the DNase-treated viromes could be attributable to a depletion of non-viral (e.g., bacterial and archaeal) DNA. Indeed, DNase-treated viromes had significantly fewer reads identifiable as 16S rRNA gene fragments by approximately two-fold (on average, 0.013% for DNase-treated, compared to 0.028% for untreated samples, Kruskal-Wallis p = 0.03, Figure 2E, Table S4). Based on taxonomic classification of these 16S rRNA gene fragments, nine of the 12 most abundant phyla across the dataset had a significantly lower abundance in the DNase-treated viromes (Figure 2F, Table S5), with Acidobacteria, Actinobacteria, and Candidatus Saccharibacteria showing the most significant differences between treatments. Bacteroidetes was the only phylum to increase in abundance in the DNase-treated viromes, and DNase treatment had no significant effect on Planctomycetes or Verrucomicrobia relative abundances.

### Viral contig and viral population (vOTU) recovery from DNase-treated compared to untreated viromes

We next wanted to assess whether treating viromes with DNase prior to DNA extraction had an influence on our ability to recover viral contigs and, subsequently, viral populations (vOTUs). We identified putative viral contigs from each single-sample assembly using VirSorter, retaining only viral contigs from the higher confidence categories (categories 1, 2, 4, and 5) (10, 18, 25). Overall, DNase-treated viromes assembled significantly more putative viral contigs (424 median, 303-475 range) in comparison to untreated viromes (226 median, 131-332 range, Figure 2A, Kruskal-Wallis P value <0.01, Table S4).

Thus far, contigs from the same viral population could have been counted multiple times (*e.g*., in assemblies from different viromes). In order to evaluate recovery of unique viral populations (vOTUs), we clustered all of the putative viral contigs from both DNase-treated and untreated viromes at 95% nucleotide identity into vOTUs (36). We then categorized these vOTUs into three groups, according to their occurrence within and/or across treatments, as follows: vOTUs containing contigs assembled from solely DNase-treated viromes (DNase vOTUs), solely from DNase untreated viromes (NoDNase vOTUs), or assembled in viromes from both treatments (shared vOTUs). In total, we identified 2,176 vOTUs, of which 1,121 were classified as DNase vOTUs, 421 as NoDNase, and 634 as shared (Figure 3, Data Set S1). Thus, DNase treatment resulted in approximately 1.7 times greater assembly of viral populations. However, of the 1,121 vOTUs that were assembled solely in DNase-treated viromes, 1,016 (90.6%) were detected in untreated viromes through read mapping, meaning that DNA from the vast majority of these vOTUs was present in the untreated viromes but did not sufficiently assemble.

**Figure 3.**
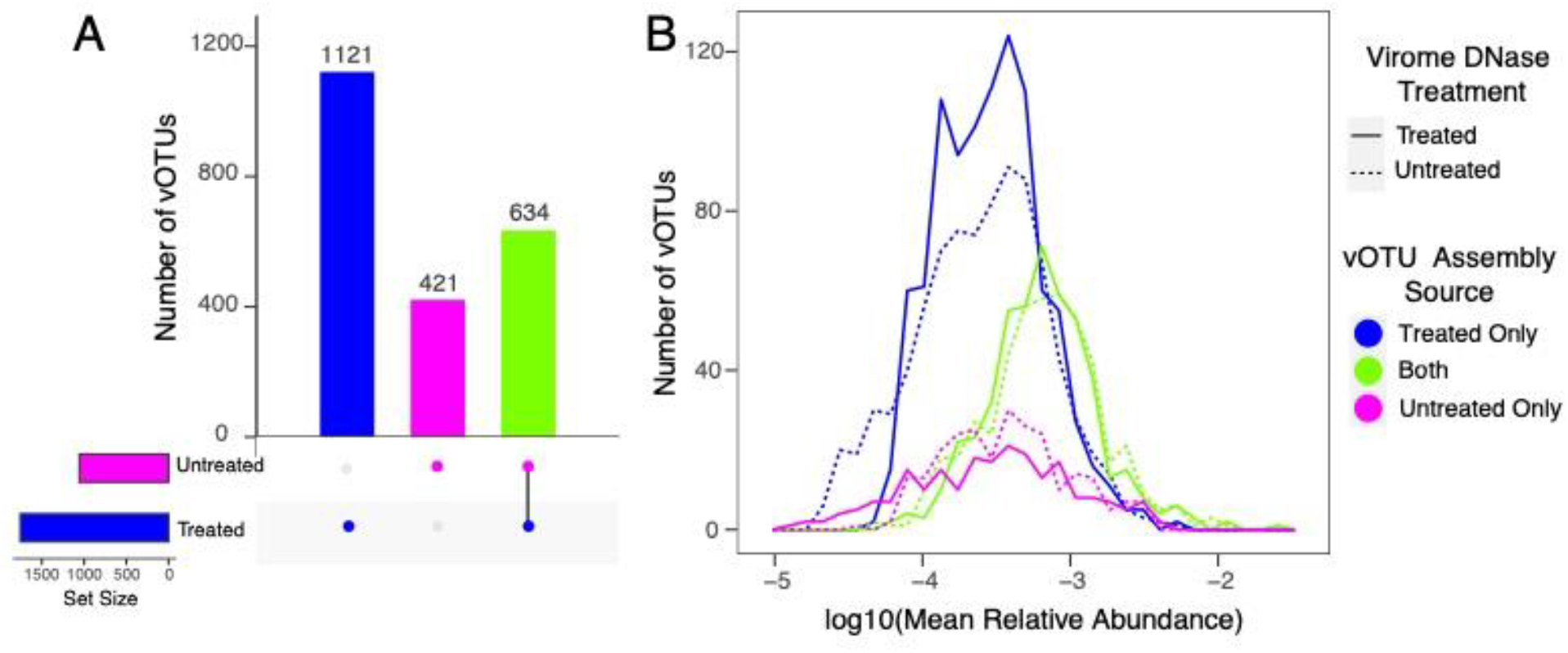
Numbers and relative abundances of vOTUs, according to the origin(s) of the assembled contigs contained within each vOTU. (A) Number of vOTUs that contained contigs clustered at 95% identity assembled from: DNase treated only (blue), untreated only (pink), or both (green) types of viromes. (B) Distribution of vOTU mean relative abundances across viromes within each DNase treatment group, colored according to the assembly source(s) of viral contigs within each vOTU. Relative abundances are derived from read mapping, such that vOTUs with contigs solely assembled from one treatment could have been detected in viromes from the other treatment via read recruitment.

### Comparing the proportions of virus-derived reads in DNase-treated and untreated viromes

With the set of 2,176 vOTUs as references for read mapping, we next sought to determine whether the smaller number of 16S rRNA gene reads was accompanied by an increase in viral reads in DNase-treated compared to untreated viromes. We mapped the quality-filtered reads from each sample to the dereplicated reference set of all vOTUs. A significantly higher number and fraction of reads from DNase-treated viromes mapped to vOTUs (on average, ~3.2 million, or 17% of reads per sample), compared to untreated viromes (on average, ~1.9 million, or 9% of reads per sample, Figure 2C, Table S4). DNase treatment improved viral enrichment more than two-fold, compared to untreated viromes.

### Patterns in the taxonomy and types of vOTUs assembled from DNase-treated and untreated viromes

We next wanted to determine whether there were differences in the types of vOTUs recovered in DNase-treated compared to untreated viromes. We performed whole-genome, network-based clustering of predicted proteins, using vConTACT2 (53), to cluster groups of vOTUs at approximately the genus level into viral clusters, or VCs (63). vConTACT2’s collection of viral genomes from the NCBI RefSeq database (ViralRefSeq-prokaryotes-v85) was included in this analysis for assigning taxonomy, as previously described (53). Of the 2,176 total vOTUs, 1,457 (67.0%) clustered into 744 VCs. 599 VCs (80.5%) contained vOTUs assembled in both treatments, while 131 VCs (17.7%) exclusively contained vOTUs assembled from DNase-treated viromes and 14 VCs (1.8%) exclusively contained vOTUs assembled from untreated viromes. Only 43 vOTUs were assigned taxonomy, based on clustering in the same VC as a reference sequence, and these vOTUs accounted for 0.3%-1.6% of the total viral community abundance (based on read mapping) in each virome. Considering these limited taxonomic assignments, DNase-treated and untreated viromes generally did recover the same taxonomic groups, namely the Caudovirales families *Siphoviridae* (10 VCs), *Myoviridae* (5 VCs), and *Podoviridae* (4 VCs). We note that these results are based on the current taxonomies for the relevant reference sequences, but phage taxonomy is actively undergoing revision by the International Committee on Taxonomy of Viruses (ICTV), and these groups of Caudovirales have been recommended for removal as taxonomic groups (64).

### Recovery of relatively rare compared to abundant vOTUs by treatment

We addressed relative abundance patterns (for example, whether recovered vOTUs tended to be relatively abundant or rare) by comparing the proportions of reads recruited to vOTUs in the three different vOTU source categories (*i.e.*, occurrence in assemblies within and/or across treatments, described earlier, Figure 4). The shared vOTUs (those assembled in at least one virome from both treatments) recruited on average 57.7% and 59.5% of mapped reads from DNase-treated and untreated viromes, respectively, despite these shared vOTUs only accounting for approximately 29% of the total vOTUs (634/2,176, Figure S2). While vOTUs uniquely assembled from DNase-treated viromes accounted for 52% of all vOTUs (1,121/2,176), they only recruited an average of 32.5% of mapped reads from DNase-treated viromes and a similar but slightly lower percentage of mapped reads from untreated viromes (28.9%) (Figure S2). These results led us to suspect that the vOTUs uniquely assembled in DNase-treated viromes tended to be relatively rare (low abundance), compared to shared vOTUs in the dataset. To address this, we constructed frequency plots of the mean relative abundances of vOTUs by category (treatment-specific or shared) (Figure 3). In both DNase-treated and untreated viromes, the distribution of treatment-specific vOTU abundances was shifted to the left (indicating lower abundances), compared to the abundances of shared vOTUs (Kruskal-Wallis p<0.001). In untreated viromes, vOTUs assembled only from untreated viromes had similar mean relative abundances to vOTUs assembled only from DNase-treated viromes. In contrast, in DNase-treated viromes, vOTUs assembled only from DNase-treated viromes had significantly higher relative abundances compared to vOTUs assembled only from untreated viromes (Kruskal-Wallis p<0.001). In short, while there were some vOTUs that were both uniquely assembled in one treatment and in high abundance in one or both treatments, the vast majority of the treatment-specific vOTUs were in low relative abundance, compared to those that were assembled in both DNase-treated and untreated viromes.

**Figure 4.**
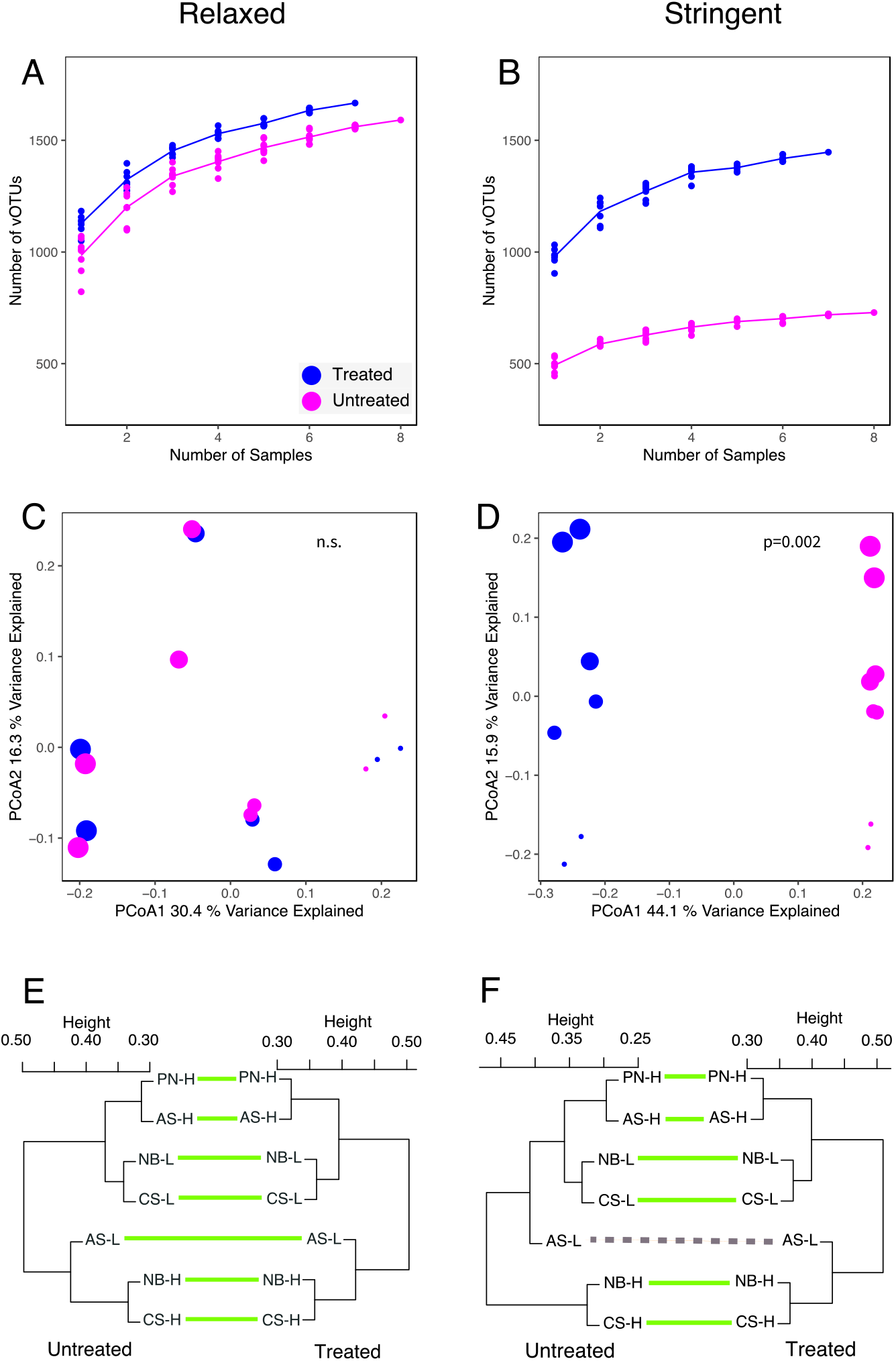
Comparisons of ecological properties across DNase treatment and vOTU detection criteria. (Left Panels) Results from the “Relaxed” vOTU detection criteria and (Right) results from the “Stringent” vOTU detection criteria (“Relaxed” detection allowed for read mapping to all vOTUs in the dataset, and “Stringent” detection allowed for mapping only to vOTUs derived from the same virome treatment group). (A&B) Accumulation curves showing the total number of vOTUs detected within a DNase treatment group at different numbers of samples. (C&D) Principal coordinates analyses of Bray-Curtis dissimilarities (each point is one virome), labeled by DNase treatment (color) and location along the E-W axis of the sampled field (shape size, largest symbols correspond to locations farthest East, with decreasing size along the E-W axis). P-values show significance of DNase treatment on community structure using PerMANOVA (E&F) Tanglegrams, each linking two sets of hierarchical clusters of viral community composition (one per DNase treatment group). Green lines connect samples with congruent clustering between the two treatment groups, and dashed lines connect samples with discongruent clustering. Numbers in the middle of each tanglegram correspond to the plot’s location along the E-W axis of the sample field. Dendrograms were created using complete linkage clustering with Bray-Curtis dissimilarities. In panels E and F, the untreated virome from plot PN-L was removed, as it did not have a paired DNase-treated virome. As a result, there was only one virome per treatment (from plot AS-L) at that particular E-W location within the field; all other E-W locations were represented by two viromes per treatment.

### Ecological inferences from DNase-treated compared to untreated viromes

In order to better understand how DNase treatment, or a lack thereof, might influence downstream ecological interpretations from soil viromic data, we applied and compared two different sets of vOTU detection criteria. For both analyses, we followed the same established best practices for considering a vOTU to be “detected” in a given sample (36). The first set of detection criteria, which we refer to as “relaxed”, considers data from reads mapped to all 2,176 reference vOTUs (i.e., vOTUs assembled from any virome in this study). The second set of criteria, referred to as “stringent”, removed from consideration reads that mapped to vOTUs that were only assembled from the other treatment group. This stringent set of criteria was meant to mimic a dataset in which only one treatment had been performed (DNase treated or untreated), as would be expected for most viromic studies. DNase-treated viromes had significantly higher perceived richness (alpha diversity) than their untreated counterparts for both the relaxed (on average 1,128 vs. 985 vOTUs, Table S6, Figure 4AB, Kruskal-Wallis p = 0.003) and stringent (on average 980 vs. 494 vOTUs per untreated, Table S6, Kruskal-Wallis p = 0.001) criteria. While both DNase-treated and untreated viromes had lower observed richness using the stringent criteria, the untreated samples showed a greater decrease (approximately two-fold) in richness between the relaxed and stringent criteria.

We also wanted to determine whether DNase treatment affected analyses of viral community structure. Using both the relaxed and stringent criteria for vOTU detection, we calculated pairwise Bray-Curtis dissimilarities between viromes. With the relaxed criteria, there was no significant difference in viral community composition attributable to DNase treatment (Figure 4C, PerMANOVA p = 0.952), but the application of the stringent criteria did result in a significant effect of DNase treatment (Figure 4D, p = 0.002). We also wanted to assess whether one set of viromes exhibited a greater amount of variation than the other. When using the relaxed vOTU criteria, the beta dispersion (i.e., the breadth of beta diversity within a group) of the DNase-treated and untreated viromes was statistically indistinguishable (Homogeneity of multivariate dispersions p = 0.430). When applying the stringent vOTU detection criteria, the DNase-treated viromes trended towards showing greater beta dispersion, but the difference was not statistically significant (Homogeneity of multivariate dispersions p = 0.075).

Finally, we previously observed a strong East-to-West gradient effect on viral community composition in these agricultural fields, using DNase-treated viromes only (30). Under the assumption that this gradient effect was real, we assessed our ability to detect this effect in both the DNase-treated and untreated viromes. We observed a significant East-West structuring of viral community composition in both sets of viromes, using both relaxed and stringent criteria (Figure 4 CD, Table S7). We further confirmed the robustness of viral community compositional patterns to DNase treatment by testing for correlations between the Bray-Curtis community dissimilarity matrices derived from DNase-treated compared to untreated viromes, using Mantel tests. The observed beta-diversity patterns (i.e., how samples were grouped according to viral community composition) were highly correlated between DNase-treated and untreated viromes, according to both stringent and relaxed vOTU detection criteria (Mantel R = 0.87 for relaxed criteria, R=0.83 for stringent criteria, both p = 0.002). This result was further reinforced in a tanglegram, which showed highly similar hierarchical clustering of samples according to viral community composition between the two virome treatments, independent of vOTU detection criteria (Figure 4 EF). A single sample (AS-L) clustered differently in tanglegrams derived from DNase-treated compared to untreated data when using the stringent vOTU detection criteria only. This was the only sample that lacked a paired sample from the same East-West position in the field, owing to the necessary removal of the single successful virome from that plot (plot PN-L) in this analysis, because its matched DNase-treated virome failed at the library construction step. Otherwise, each pair of samples from the same field column grouped together in all four hierarchical clusters (DNase-treated vs. untreated and relaxed vs. stringent vOTU detection criteria).

### Comparing viral recovery from untreated viromes and total soil metagenomes

In most metrics that we have compared to this point, DNase-treated viromes have outperformed untreated viromes, but we wanted to know the extent to which untreated viromes could still improve viral sequence recovery and reduce bacterial and archaeal DNA content in viromes, compared to total soil metagenomes. We previously analyzed total soil metagenomes from these same samples (30), which showed an average of 2.2% viral contig content (compared to 45% for untreated viromes in this study, a ~20-fold improvement), 0.04% 16S rRNA gene reads (compared to 0.02% for untreated viromes here), and an average of 0.04% of reads mapping to vOTUs (compared to 9.2% for untreated viromes here, a ~225-fold improvement). Further, the ecological patterns observed in this study were robust to different DNase treatments (Figure 4), and we wanted to know the extent to which mining total metagenomes for viral signatures would yield the same patterns. For example, a highly significant effect of spatial structuring (E-W gradient effect) on viral community composition was observed for untreated viromes here, even with the stringent detection criteria (PerMANOVA p=0.003), and we wanted to know the extent to which this pattern could also have been recovered from the total soil metagenomes. While this result was reproduced with viral communities recovered from the total soil metagenomes, the significance was borderline (PerMANOVA p=0.045).

## Discussion

### DNase treatment of viromes reduced contamination and sequence complexity, consistent with removal of free DNA

We have shown that DNase treatment of viromes significantly reduced sequence complexity and lowered the amount of contaminating cell-derived DNA (measured as 16S rRNA gene fragments) by about two-fold. Sequence complexity has long been a challenge for assembling environmental metagenomes and can result in high fragmentation of genomes from low-abundance species (65, 66). Thus, we suspect that the observed decrease in sequence complexity in DNase-treated viromes was responsible for the larger, more contiguous assemblies from DNase-treated viromes, and it is reasonable to assume that this reduction in sequence complexity resulted from free (“relic”) DNA depletion as a result of successful DNase treatment.

Relic DNA (sometimes called environmental DNA, eDNA, or free DNA) is not contained within a viable cell or virion and has been shown to artificially increase the observed richness of microbial communities in some soils (38, 39), presumably by allowing for the detection of locally dead or extinct microbial taxa (38). Studies have also suggested that the presence of relic DNA can obscure or minimize patterns in beta diversity (38, 67, 68), but here we observed that both DNase-treated and untreated viromes produced viral communities with highly correlated beta-diversity patterns (Figure 4). Although there was a single sample that clustered differently in DNase-treated compared to untreated viromes when using the stringent vOTU detection criteria, we attribute this difference predominantly to the lack of a successful replicate matching sample in the same column of the field, rather than differences in relic DNA composition between treatments.

### Viromics without DNase treatment might be particularly useful for samples stored frozen

The laboratory protocol for generating viromes requires equipment that is unlikely to be available or practical to run in the field, precluding immediate processing of samples collected from distant field sites (19, 34). Even samples from nearby sites may need to be stored temporarily, as a relatively small number of samples can be processed for viromics at a time (6-12 per ~2 days in our lab, but this will depend on available equipment and personnel). Frozen storage can preserve *in situ* community composition (69–72), and ideally, virions would be frozen in cryoprotectant or similar to preserve their integrity, but the compatibility of cryoprotectants with various viromics protocols is not well known. Thus, in some cases, direct freezing of samples may be necessary (19). We have previously shown that freezing can prohibit the use of DNase on aquatic viromes, resulting in viral DNA yields below detection limits (34), and anecdotally, we see have seen similar results from soils stored frozen (data not shown).

Encouragingly, work from our group has shown that viromes prepared without DNase treatment (untreated viromes) from frozen peat soils can still substantially improve vOTU recovery, compared to total metagenomes (19). Similarly, hypersaline lake water stored frozen yielded predominantly viral sequences in viromes that did not undergo DNase treatment (34). In combination with the complete depletion of DNA after DNase treatment in these hypersaline lake samples, it is reasonable to suspect that some virions became compromised by freezing, such that DNase treatment removed valuable viral genomic DNA contained in degraded virions that may have been intact in the field. Studies in pure culture support virion degradation through freezing; for example, coliphages from wasterwater showed decreased viability after prolonged storage in frozen wastewater, and *Bacillus subtilis* bacteriophage viability decreased multiple logs after only two hours of frozen storage with no cryoprotectant (43, 44).

Here, to ensure that we could get sufficient DNA for sequencing from both treatments for a direct comparison, we compared fresh samples with and without DNase treatment. While results from this and prior studies converge to suggest that skipping DNase treatment is likely to be a good option for viromics from samples stored frozen, future comparisons would benefit from including a combination of samples processed fresh and after frozen storage.

### Recommendations for future viral ecology studies

We have shown here that DNase treatment produced better assemblies, more viral contigs, fewer 16S rRNA gene reads (indicative of bacterial and archaeal DNA), and more viral reads in comparison to not treating with DNase. However, both kinds of viromes substantially outperformed total soil metagenomes in these metrics. Together, these results suggest that soil viromics with DNase treatment is the best approach for interrogating soil viral ecology, where possible, but soil virome preparation without DNase treatment can be better than total soil metagenomics when DNase treatment is not an option. Prior work suggests that these results may be generalizable to viromics in other ecosystems as well (34), but to our knowledge, direct comparisons of these approaches have not been made in other ecosystems. The decision of what approach to take is inherently dependent on the questions being asked, along with the logistics of sample collection, storage (and possibly shipment), and performing the laboratory sample processing. Where possible, we recommend processing soil viromes soon after sampling and including a DNase treatment after virion purification and prior to virion lysis.

However, even without DNase, soil viromes substantially enrich for viral sequences in comparison to total soil metagenomes. Across multiple studies, including fresh, frozen, agricultural, and peat soils in various combinations (this study and (19, 30)), viromics (with or without DNase) seems to outperform total metagenomics for soil viral community investigations. Still, only a tiny fraction of soil types and a few combinations of laboratory procedures have been attempted, so assessing the broad generalizability of the observed trends will require expanding our investigations across diverse terrestrial and other ecosystems. Thus far, the extra effort required to purify virions from soil prior to DNA extraction seems to be worthwhile, even without DNase treatment.

## Acknowledgements

We thank Sanjai Parikh and Danielle Gelardi for designing and maintaining the field experiment from which all samples for this study were derived. We thank Sara Geonczy, Winston Bess, and Rose Bolle for contributing to discussions of this work.

This work was supported by the UC Davis College of Agricultural and Environmental Sciences and Department of Plant Pathology (new lab start-up to JBE). This work was also supported in part by the USDA National Institute of Food and Agriculture, Hatch project number CA-D-PPA-2464-H/accession number 1016718. CSM was supported by an award from the U.S. Department of Energy, Office of Science, Office of Biological and Environmental Research, Genomic Science Program, #DE-SC0020163 (grant to JBE). The sequencing was performed by the UC Davis Genome Center’s DNA Technologies and Expression Analysis Core, supported by NIH Shared Instrumentation Grant 1S10OD010786-01. Any opinions, findings, conclusions, or recommendations expressed in this manuscript are those of the authors and do not necessarily reflect the view(s) of any funding agencies or institutions.

